# Smartphone behavior is suppressed during bursts of wake slow waves

**DOI:** 10.1101/2025.10.06.680725

**Authors:** Ruchella Kock, David Hof, Reto Huber, Arko Ghosh

## Abstract

Real-world behavior unfolds as a continuous flow of actions over time. One possibility is that this continuous flow is enabled by a seamless stream of cortical activity. However, emerging evidence suggests that even during ongoing behaviors, cortical neural populations undergo brief periods of silence. This ‘off-state’ can be detected as wake slow waves in EEG, similar to the slow waves characterizing deep sleep. Here, we leveraged the spontaneously occurring fluctuations in smartphone touchscreen interactions in combination with EEG to investigate the dynamics of slow waves during behavior. We find that slow waves occur in bursts - characterized by short consecutive intervals - interleaved with longer periods of absent slow waves. During smartphone use, the probability of bursts increased compared to rest, and the duration of burst-absent periods was lower. During the bursts, however, the likelihood of generating smartphone interactions was substantially diminished. Conversely, the periods of absent slow waves were more permissive to the behavioral output. We propose that slow wave burst dynamics are state-dependent, configured to create temporal windows permissive to action generation. This configuration, in turn, allows behavioral outputs to leverage the windows of opportunity that emerge between slow wave bursts. Our study, combining smartphone behavior with EEG, reveals how brief cortical silencing events structure the computations underlying everyday actions.

## 1 Introduction

Everyday behavior unfolds as a continuous stream of actions - typing, reaching, scrolling - with no apparent gaps. Theories of cortical function often assume that cortical activity may seamlessly continue without any breaks [Histed et al., 2009, Huk et al., 2018]. For instance, according to invasive recordings in non-human primates, prefrontal cortical activity can seamlessly persist from one task to the next, and the cortical preparation for the subsequent tasks can begin well before the completion of the current task [Histed et al., 2009]. However, research on cortical dynamics during sleep reveals that cortical neurons can undergo periods of silence (off-state) followed by high neural firing (on-state) [Steriade et al., 1993]. These off-states can break the continuous stream of cortical processing and disconnect cortical networks [Massimini et al., 2005]. While these off-states are a defining feature of deep sleep, growing evidence shows that they can also occur during wakefulness, albeit more localized than in sleep [Bernardi et al., 2015, Vyazovskiy et al., 2011, Nir et al., 2017, Hung et al., 2013]. In EEG recordings, these cortical off-states are captured as large-amplitude slow waves in the delta (1–4 Hz) or theta (4–7 Hz) range [Sachdev et al., 2015, Andrillon et al., 2021]. Although the occurrence of wake slow waves is increasingly studied for isolated behavioral tasks [Andrillon et al., 2021], whether these cortical silencing events constrain behaviour during naturalistic, ongoing actions remains unknown.

Traditional homeostatic models emphasize time-dependent sleep pressure accumulation, low-frequency activity (including slow waves) increases with time awake, but do not account for how different behavioral states might modulate slow wave dynamics [Borbély et al., 1982]. Supporting this idea, sleep deprivation studies show that wake slow waves become more frequent and higher in amplitude as the time spent awake is experimentally increased [Finelli et al., 2000, Snipes et al., 2023]. In contrast, other research suggests that the distribution of slow waves is shaped by other factors, particularly by cognitive demand and local plasticity processes [Halász et al., 2014, Huber et al., 2004, Huber et al., 2006, Snipes et al., 2022, Hung et al., 2013, Massimini et al., 2005, Quercia et al., 2018, Pinggal et al., 2022]. While these perspectives establish that slow waves are modulated at multiple timescales, from hours-long homeostatic processes to state-dependent fluctuations, how slow wave dynamics relate to moment-to-moment behavioral output in naturalistic contexts remains unexplored.

The intervals between slow waves may be critical for understanding their state-dependent operations, as these intervals could reflect cortical states that are more or less conducive to behaviour. This idea stems from cortical stimulation studies, where experimentally induced neural interruptions at different intervals provide an analog for naturally occurring slow wave events [Cash et al., 2010, Benali et al., 2011, Ferreri et al., 2011, Bolognini and Ro, 2010]. This relationship is nonlinear: performance effects on artificial tasks emerge only within specific interval ranges [Luber et al., 2007] and depending on timing, induced interruptions can paradoxically improve performance on some tasks [Luber and Lisanby, 2014]. However, while sleep slow waves exhibit nonlinear temporal dynamics [Wei et al., 2020, Chauvette et al., 2011], the interval statistics of wake slow waves during ongoing behaviour remain uncharacterized.

A powerful approach for characterizing nonlinear systems is to examine the intervals between successive events and how one interval predicts the next. This approach is inspired by Poincaré sections, where continuous high-dimensional dynamics can be understood through discrete events sampled at a lower-dimensional cross-section. It has been extensively used to quantify spiking activity, and cardiac, and behavioural systems, and here we apply it to characterize the temporal organization of wake slow waves during naturalistic behaviour [Brennan et al., 2002, Rodieck et al., 1962, Eagan and Partridge, 1989, Ceolini and Ghosh, 2023, Duckrow et al., 2021]. Essentially, the interval between two consecutive wake slow waves (K) can be considered along with the next interval (K+1). These intervals can then be pooled in two-dimensional bins by calculating the joint distribution. These bins can capture the probability of slow waves, separating rapid (short intervals, say 300 ms) from isolated slow waves (long intervals, say 30 s). By applying this analysis to EEG-detected slow waves, we can test whether their temporal organization relates to the moment-to-moment behavioral fluctuations. Traditional laboratory tasks are too constrained to capture naturalistic neuro-behavioral dynamics.

Smartphone use engages a range of cognitive processes, provides fine-grained quantitative data, and is an emerging approach to unravel complex neuro-behavioral links [Wan and Ghosh, 2025, van de Ruit and Ghosh, 2022, Duckrow et al., 2021, Kock et al., 2023, Balerna and Ghosh, 2018, Gindrat et al., 2015]. Here we quantified the distribution of slow waves during ∼ 60 minutes of smartphone use. First, we compared slow wave properties during smartphone use to a passive movie viewing baseline [Meer et al., 2020, Espenhahn et al., 2020]—chosen to sustain a wakeful, low-demand state for 60 minutes—revealing state-dependent differences at hourly timescales. Next, we characterized the fine-grained temporal organization of slow waves using joint-interval distributions, revealing distinct burst patterns between sessions. Finally, using high-resolution smartphone touchscreen logs, we found that touchscreen interaction rates were reduced during periods when slow waves occurred in bursts. These findings reveal that nonlinear slow wave dynamics structure real-world behavior.

## 2 Results

### Slow waves in smartphone behavioral session and rest

We recorded EEG from 46 participants during a session of passive movie watching (∼60 minutes) to establish a baseline resting state, followed by a session of smartphone use (∼60 minutes). The recording sessions were distributed from morning to late afternoon, with session onset times ranging from 7 am to 5 pm (Supplementary Figure 1). We monitored sleep patterns during the preceding week (Supplementary Figure 2) to ensure participants were not sleep-deprived. Based on homeostatic models of slow wave regulation, minimal accumulation is expected over our hour-long observation window in well-rested participants [Borbély et al., 1982]. To confirm this, we fitted a linear regression model predicting slow wave density per minute (number of slow waves within 1-min time windows) from session type (movie vs. smartphone), time elapsed within session, and their interaction (Supplementary Figure 3). Neither time elapsed nor the interaction term reached statistical significance; likewise, time elapsed was not significant in a reduced model without the interaction term (all *p >* 0.05 after multiple comparison correction, MCC). Session type, however, significantly predicted slow wave density, with smartphone use showing higher density than movie watching (detailed results below and in equivalent direct comparison of slow wave density between sessions). These results confirm that homeostatic slow wave accumulation was negligible over our observation period, validating subsequent analyses of state-dependent dynamics.

To characterize the state-dependent differences, we quantified three key features of slow waves: (i) density (frequency of neural silencing), (ii) amplitude (extent of neural synchrony), and (iii) upwards and downwards slopes (indices of synaptic strength, with steeper slopes reflecting stronger synapses [Jaramillo et al., 2020]). We performed mass univariate statistics using each of these features, comparing the two sessions to reveal spatial clusters of significant differences (p ¡ 0.05, MCC using cluster-based permutation correction). Slow waves were present across the scalp in both sessions. At rest, slow wave topography replicated prior findings [Andrillon et al., 2021]. Specifically, density was highest at central electrodes, amplitude was largest at frontal sites, and slopes were shallowest in sensorimotor areas. During smartphone behavior, the density of slow waves was higher compared to rest, except for the occipital areas (Figure 1). Amplitudes were reduced during smartphone behavior at frontal and temporal sites, and remained unchanged at the occipital electrodes. The slopes were shallower than at rest across most of the scalp, except at central and occipital regions, where they were steeper. The increased density but reduced amplitude of slow waves during smartphone use, occurring without homeostatic accumulation, indicates state-dependent modulation in both the frequency and synchrony of cortical silencing events.

**Figure 1.**
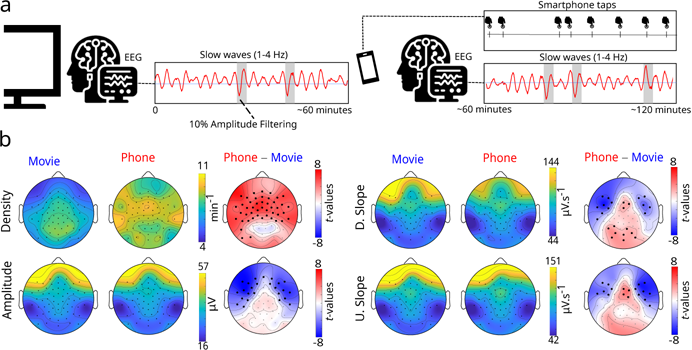
Slow waves were observed during passive movie watching and smartphone use. **(a)** Schematic overview of the experimental protocol: participants underwent ∼60 minutes of restful movie watching (rest) followed by ∼60 minutes of smartphone use, during which slow waves were captured with EEG. **(b)** We contrasted slow wave features between smartphone use and rest using mass-univariate paired t-tests. Scalp topographies display population-average slow wave features: density (slow waves per minute), amplitude, and downward/upward slopes. For each feature, the left topography shows the mean during rest, the middle topography shows the mean during smartphone use, and the right topography shows *t* -values from the paired t-tests. Statistical significance was assessed with multiple comparison correction in the form of cluster-based permutation testing (*α* = 0.05; 1000 randomizations). Electrodes showing statistically significant differences are marked with black dots on the topographical *t* -value maps. Slow wave density was higher during smartphone use across most of the scalp, with no significant difference at parieto-occipital electrodes. In contrast, amplitude was larger at rest, particularly over frontotemporal regions. Slopes (upward slope, U, and downward slope, D) were steeper at rest in frontotemporal areas, but shallower in central and occipital regions.

A potential concern is whether touchscreen interactions evoke event-related potentials (ERPs) misclassified as slow waves. Three lines of evidence argue against this: First, slow waves were observed during rest when no touchscreen interactions occurred. Second, smartphone-related ERPs differ markedly from slow waves in amplitude ( 1 µV vs 10 µV). Third, these ERPs are localized to contralateral sensorimotor cortex whereas slow waves were detected bilaterally across the entire scalp [Kock et al., 2023].

### Minute-scale relationships between slow waves and fluctuations in smartphone interactions

To examine whether slow waves relate to behavioral fluctuations at minute-scale resolution, we binned EEG and behavioral data into 1-minute windows [van de Ruit and Ghosh, 2022]. This approach tests for monotonic relationships between slow wave properties and behavioral output at coarse temporal scales. We computed the slow wave density and median amplitudes (note we dropped slopes as a feature due to its redundancy with amplitudes) and correlated these with the number of smartphone interactions occurring in the same bins. First, we performed separate Spearman correlations between interaction counts and each slow wave feature (density and amplitude) at each electrode (Supplementary Figure 4). Isolated electrodes showed correlations in individual participants, but no consistent patterns emerged at the group level. Next, we conducted a hierarchical mass-univariate analysis predicting interaction counts from slow wave density and amplitude at each electrode (within subjects), followed by group-level one-sample t-tests on the resulting regression coefficients. We found significant clusters over the frontotemporal and sensorimotor regions where density was negatively associated with interaction rates (Supplementary Figure 4). However, the slow wave amplitude did not reveal significant clusters. Interestingly, although overall slow wave density was higher during smartphone use (see above), moment-to-moment increases in density within the smartphone session were associated with decreased interaction rates, suggesting that slow wave bursts may transiently suppress behavior.

### Fine-grained dynamics of slow waves in smartphone behavioral sessions and rest

Having described associations at minute-scale resolution, we next examined the fine-grained temporal organization of slow waves by analyzing intervals between consecutive slow waves at each electrode. Specifically, we captured the probability of observing a given interval duration followed by a specific subsequent interval duration, forming a two-dimensional joint-interval distribution spanning from 30 ms to 30 s (JID; Figure 2, Supplementary Figure 5). This approach, based on Poincaré sections that reduce continuous dynamics to discrete event sequences (see Introduction), examines how one inter-event interval follows the next, revealing whether slow waves occur randomly, in bursts, or with other temporal structure. We performed two complementary analyses of these distributions: First, we used non-negative matrix factorization (NNMF) to identify prototypical temporal patterns shared across electrodes in movie watching and smartphone use sessions. Second, we tested for session differences at each point in the JID using mass-univariate paired t-tests.

**Figure 2.**
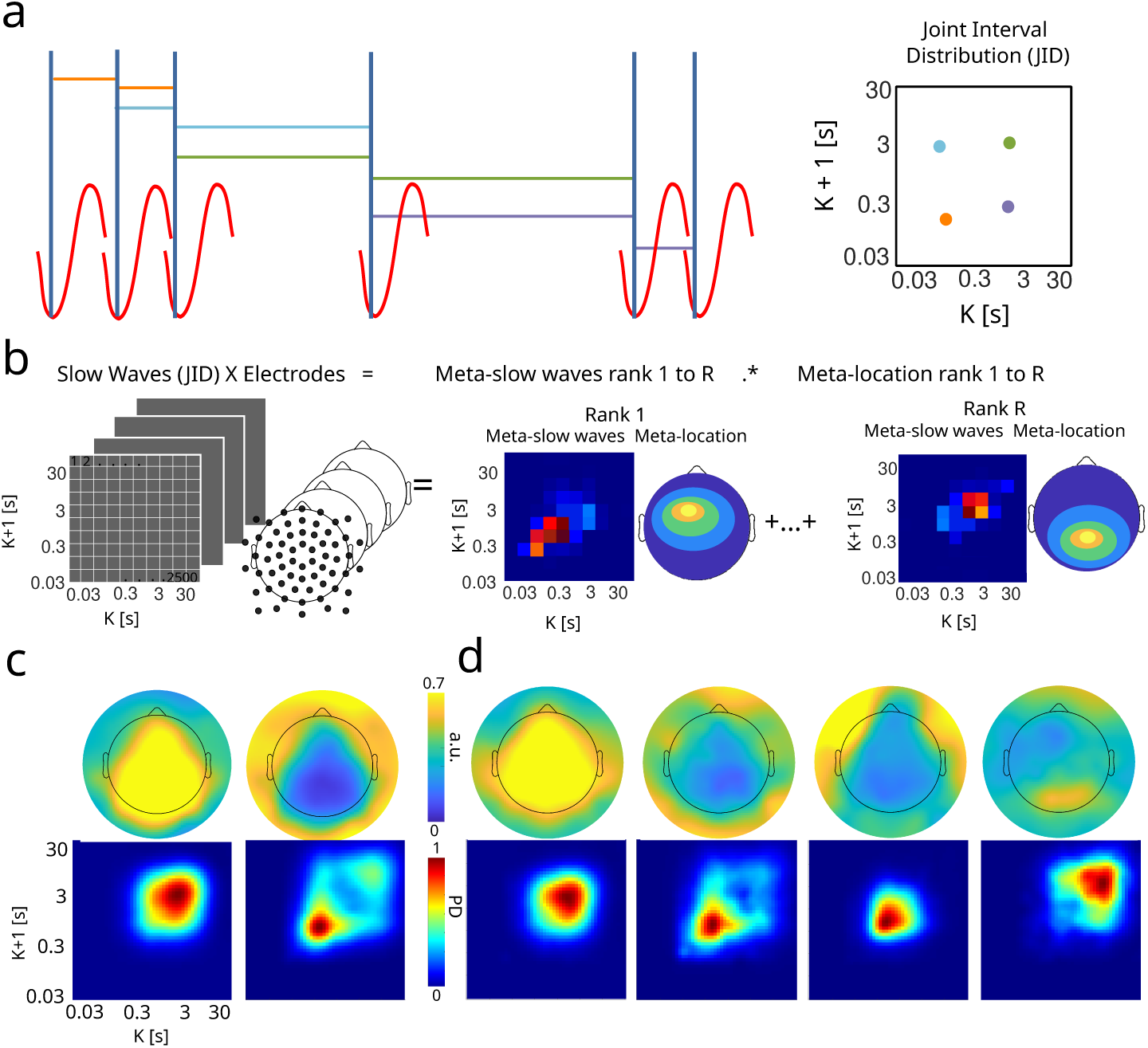
Prototypical fine-grained slow wave temporal dynamics identified across the population. **(a)** Method schematic describes how the interval between two consecutive slow waves (K) can be considered along with the next interval (K+1). The probabilities of the intervals are pooled in two-dimensional bins, resulting in the joint-interval distribution (JID). **(b)** Schematic showing how the temporal dynamics of slow waves and their spatial locations can be derived using non-negative matrix factorization (NNMF). For each participant, NNMF decomposes a matrix of slow wave JIDs per electrode into low-rank representations: meta-JIDs capturing the typical slow wave dynamics, and meta-locations indicating where these dynamics typically occur on the scalp. **(c)** We clustered the meta-JIDs derived at rest across the population using k-means, and display the population-level cluster centroids. Scalp topographies show the average (20% trimmed mean) meta-location for each cluster. We identified two clusters: (i) long consecutive intervals at central, parietal, and occipital electrodes, and (ii) rapid consecutive slow wave intervals (bursts) in the frontal areas. The dynamics are shown in *log*_10_ space. **(d)** The same analysis was performed for smartphone use, where four clusters were identified: (i) long consecutive intervals at central, parietal, and occipital electrodes, resembling rest; (ii) bursts irregularly distributed across the scalp; (iii) moderate bursts concentrated in the frontal areas; and (iv) long consecutive intervals localized to occipital electrodes.

NNMF decomposed the high-dimensional JID × electrode tensor into pairs of basis vectors, each comprising a prototypical interval pattern (which we term a ‘meta-JID’) and its corresponding spatial distribution across electrodes (‘meta-location’). Importantly, both sessions showed two dominant clusters per individual (movie watching median=2, range= 1 - 3, and smartphone use median=2, range= 1 - 6). Clustering these patterns at the population level revealed two dominant classes at rest: (r.i) long consecutive slow wave intervals, occurring primarily at central, parietal, and occipital electrodes, and (r.ii) rapid consecutive slow wave intervals (i.e., bursts), largely absent from central, parietal, and occipital regions but prevalent at frontal electrodes. During smartphone behavior, four dominant classes emerged: (s.i) a pattern similar to that seen at rest, with long consecutive slow wave intervals across central, parietal, and occipital electrodes; (s.ii) also similar to rest, bursts, but less prominent over central electrodes and more irregularly distributed across the scalp than at rest; and unique to smartphone use: (s.iii) moderate bursts concentrated at the frontal electrodes; and (s.iv) long consecutive slow wave intervals localized to occipital electrodes. These findings reveal both shared and distinct slow wave dynamics: long-interval slow waves at central-posterior sites occur in both conditions (r.i, s.i), while smartphone use additionally engages frontal burst dynamics (s.ii, s.iii) and occipital-localized long-interval patterns (s.iv).

Next, we performed mass univariate paired t-tests spanning the scalp electrodes and the two-dimensional bins of the JID to compare the slow wave probability between rest and smartphone behavior (Figure 3, select electrodes, and Supplementary Figure 6, all electrodes). This revealed higher probability in the short-interval region of the JID (indicating bursts) during smartphone use and lower probability in the long-interval region (isolated slow waves). Statistically significant JID clusters were observed across the scalp, except at the occipital electrodes. Both analyses converge on a key finding: smartphone use increases burst probability across frontocentral and temporal regions compared to rest, with occipital electrodes showing distinct dynamics. These results demonstrate that the fine-grained temporal organization of slow waves is state-dependent, reflecting the engagement of different cognitive and motor processes during smartphone interaction.

**Figure 3.**
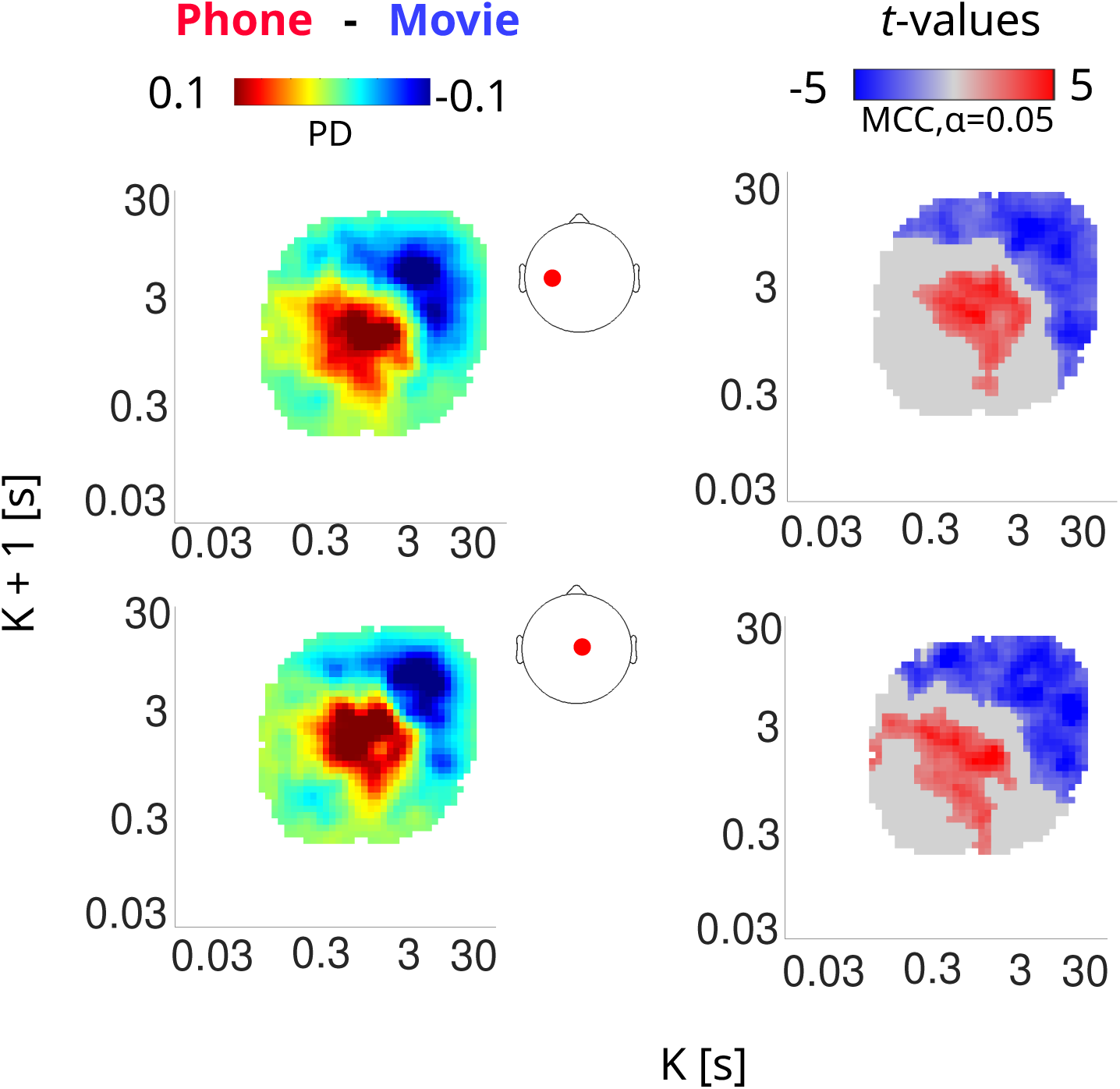
Comparison of fine-grained slow wave dynamics between rest and smartphone use. Here we show slow wave joint interval distributions (JID) from representative electrodes (red dots) at the central and sensorimotor areas. Paired t-tests showed that smartphone use was accompanied by rapid bursts of slow waves, while at rest, slow waves appeared more isolated. Mean differences between slow waves JIDs (smartphone behavior - restful movie watching) are shown in the left column. Statistically significant *t* -values are displayed in the right column. Statistical significance was assessed using spatiotemporal clustering with multiple comparison correction (MCC, *α* = 0.05, 1000 bootstraps). Grey indicates no statistically significant clusters and white areas indicate bins with no detected events. PD = probability density. For all electrodes, see the Supplementary Figure 6.

### Smartphone taps are suppressed during slow wave bursts

Both minute-scale and fine-grained interval analyses revealed associations between slow wave dynamics and smartphone behavior, with higher slow wave density related to reduced interaction rates. Given the distinct patterns associated with bursts and isolated slow waves, we next report on the nature of the behavioral output during these different dynamics. To examine this relationship, we estimated the rate of smartphone touchscreen interactions as a function of the slow wave dynamics captured using the JID, at each electrode. For each pair of consecutive slow wave intervals (K, K+1), we calculated the number of interactions occurring between the corresponding three slow waves, normalized by the total interval duration (K + K+1) (Figure 4, Supplementary Figure 7). Upon population-level averaging, we consistently observed a pattern across all electrodes: the rate of smartphone interactions was lower (∼0) during slow wave bursts, and the rate increased from short consecutive to long consecutive interval combinations (along the JID diagonal). We next addressed the latency to smartphone interactions after the slow wave, as a function of the JID (Supplementary Figure 8). We found a marginal gradient: the latency was longest (∼54 s) following rapid consecutive slow waves, and gradually shortened (∼1 s) as slow waves became sparser. These findings demonstrate that slow wave bursts are associated with transiently suppressed behavioral output: during rapid consecutive slow waves (bursts), interaction rates drop to near zero, recovering as intervals lengthen. This coupling between slow wave temporal structure and behavioral output was consistent across frontocentral and temporal regions, revealing that cortical silencing events may constrain when actions can be generated.

**Figure 4.**
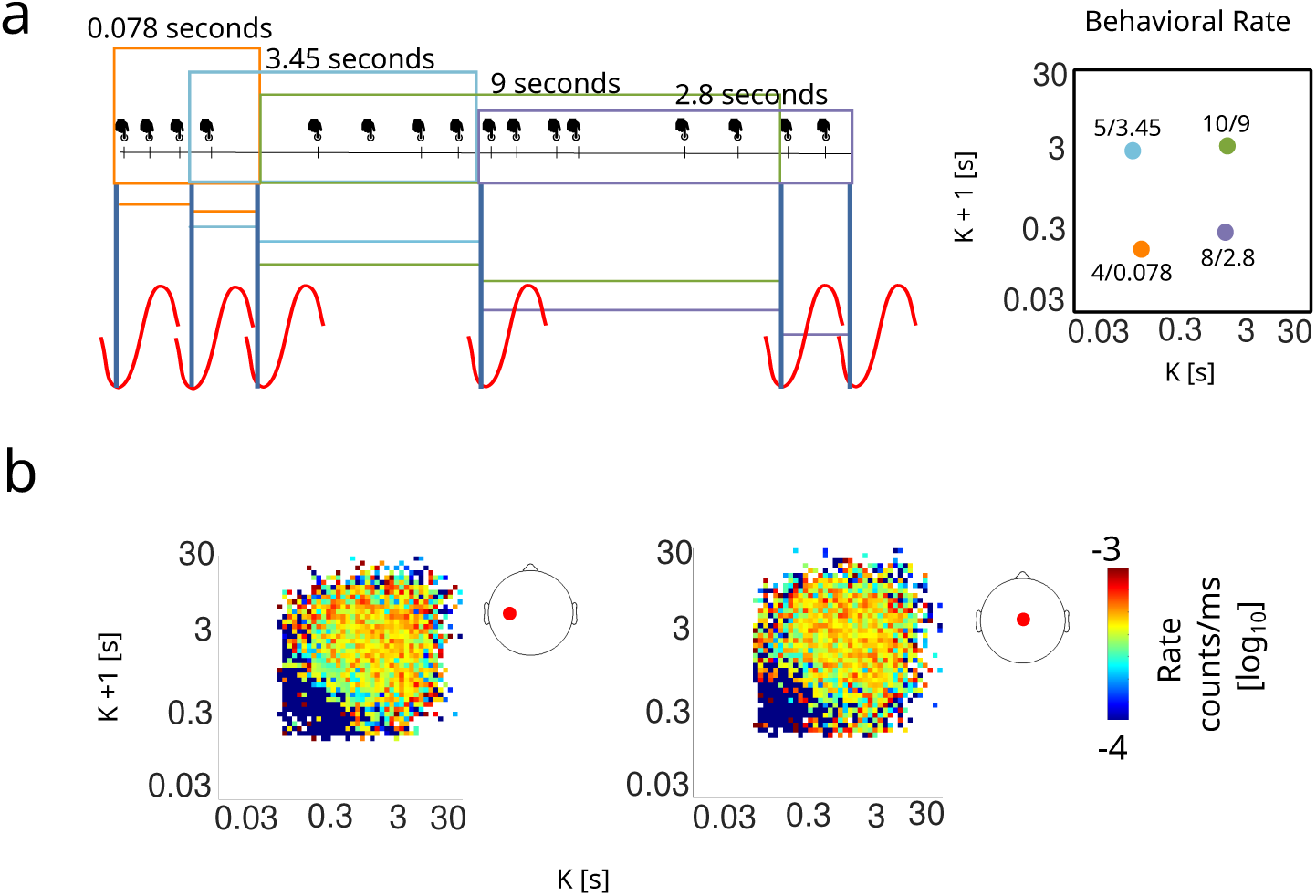
Smartphone touchscreen interactions are suppressed during slow wave bursts. **(a)** Schematic illustration of how the rate of smartphone interactions was calculated as the number of interactions occurring between consecutive slow waves (during intervals K and K+1), normalized by the total interval duration (K + K+1). **(b)** Population-level interaction rates computed using median across participants (N = 46). Representative electrodes from central and sensorimotor regions (locations marked with red dots on scalp inset) showed a systematic gradient: interaction rates were near zero during short consecutive intervals (bursts) and progressively increased toward long consecutive intervals. White areas indicate bins with no slow wave pairs. For all electrodes, see the Supplementary Figure 7.

To further characterize the distinction between bursts and isolated slow waves, we examined how amplitude (Supplementary Figure 9) and duration (Supplementary Figure 10) varied with the interval structure. In particular, we examined how slow wave features varied as a function of their temporal context (position in the JID). The median amplitude (averaged across participants) differed between bursts (short consecutive) and isolated events (long consecutive): bursts were associated with larger amplitudes relative to the isolated slow waves. This pattern was visible across the scalp. The median slow wave duration (averaged across participants) also varied along the diagonal of the JID, with short-lasting slow waves during bursts (∼500 ms). These findings demonstrate that bursts and isolated slow waves reflect distinct neural states: bursts involve briefer but more synchronized cortical silencing (larger amplitude, shorter duration) that suppresses behavior, while isolated slow waves involve longer, less synchronized off-states that permit behavioral output.

## 3 Discussion

Our findings reveal that cortical slow waves are not simply markers of drowsiness but are actively modulated by behavioral state. We compared slow waves during smartphone use and rest at two timescales: hourly (overall density) and millisecond-to-second (interval dynamics). Both analyses revealed state-dependent differences and that the temporal distribution of slow waves differs between the two sessions. The density of slow waves was higher in the smartphone use session. The fine-grained analysis revealed a spectrum of slow wave dynamics, from rapid consecutive intervals (i.e., bursts) to long consecutive intervals (isolated slow waves). Although smartphone use increased burst frequency, individual bursts suppressed behavioral output, creating a paradox: more frequent silencing during active behavior. This apparent paradox resolves when considering the multi-scale organization of slow wave dynamics. At coarse timescales (minutes), overall behavioral state (smartphone use vs. rest) modulates slow wave density. At fine timescales (milliseconds to seconds), bursts suppress behavioral output while burst-free intervals permit action generation, creating temporal windows for behavior.

Our findings confirm that homeostatic accumulation is not the only driver of slow wave dynamics at short timescales. Several lines of evidence support this conclusion. First, slow wave density showed no linear increase over time within either session. Second, interval dynamics (captured in JIDs) showed no signatures of progressive changes, such as systematically shortening intervals. Third, slow waves tracked moment-to-moment behavioral fluctuations within the smartphone session, a pattern more consistent with state-dependent modulation than gradual accumulation. However, our participants were well-rested, and whether these behavioral correlates persist under conditions of high sleep pressure remains to be determined. Sleep deprivation might alter the balance between state-dependent and homeostatic modulation of slow wave dynamics. Slow wave dynamics varied systematically with behavioral state: smartphone use increased density and burst frequency compared to rest. At minute-scale resolution, periods with more interactions showed higher overall slow wave density, though within-minute bursts suppressed individual interactions. This suggests a dynamic, state-dependent modulation of slow wave properties.

These findings raise questions about the functional significance of burst dynamics. Slow wave bursts were interleaved with long burst-free periods, even at rest. These intervals may provide windows for cortical processing, integration, and potentially network processing [Hinard et al., 2012, Krueger and Roy, 2016], even in the absence of overt behavior. The suppressive effects of bursts persisted beyond the slow waves themselves, as evidenced by prolonged latencies to subsequent interactions. This lingering suppression was specific to bursts, not isolated slow waves. The stronger behavioral suppression during bursts may reflect their larger amplitudes, which could indicate more extensive or synchronized cortical silencing. The transient behavioral suppression likely reflects a more general gating mechanism affecting cortical computations. This mechanism operates during both rest and active behavior—bursts occurred in both states, though their behavioral consequences were observable only during smartphone use. Thus, continuous behavior does not require uninterrupted cortical processing; rather, behavioral outputs are strategically timed to occur in the windows between recurrent silencing events.

Beyond their role in transiently gating behavioral output, the functional significance of wake slow waves remains unclear [Massimini et al., 2024]. One speculative possibility, based on animal studies, is that these slow waves locally enhance responsiveness to incoming stimuli [Crochet et al., 2005, Vyazovskiy et al., 2013]. Bursts may thus support periods of enhanced sensitivity to incoming information, while burst-free periods permit endogenous processing and action generation. The active engagement with tactile and visual information during smartphone use may require more frequent periods of heightened sensory sensitivity, potentially explaining the increased burst frequency. However, if bursts enhance sensory responsiveness, why do they suppress motor output? One possibility is that sensory enhancement and motor suppression could be dissociable, with slow waves creating states optimal for sensory processing but suboptimal for motor execution. An entirely different possibility is that wake slow waves serve restorative rather than computational functions akin to sleep slow waves, which have been associated with restorative processes, including synaptic homeostasis [Tononi and Cirelli, 2006], metabolic waste clearance [Xie et al., 2013], and memory consolidation [Diekelmann and Born, 2010]. These computational and restorative hypotheses are not mutually exclusive—slow waves may serve different functions at different timescales.

Transient periods of neural silencing punctuate smartphone interactions, with behavioral output tightly coupled to these dynamics. Our findings address a critical question: how does behavior flow continuously despite recurrent neural silencing? Behavioral outputs are timed to occur in burst-free intervals. Smartphone use increased burst frequency compared to rest, yet individual bursts suppressed interactions—a paradox resolved by recognizing multi-scale temporal organization. Understanding these dynamics has practical implications: after sleep loss, when homeostatic sleep pressure increases, this gating mechanism may become compromised, increasing human error [Nir et al., 2017]. These findings establish wake slow waves as active, state-dependent regulators of behavior, expanding slow wave research beyond sleep to reveal how sub-second cortical dynamics structure real-world actions.

## 4 Methods

### Participants

Fifty-two (52) healthy participants ranging between 23 to 52 years old (median = 29) were recruited on Leiden University campus through flyer-based advertisements. The inclusion criteria were being right-handed (self-report), free of neurological disorder and medication, and unshared ownership of an Android operating smartphone. The procedures were approved by Leiden University Psychology Research Ethics Committee, Leiden, The Netherlands (Number: V1-920) on 07-04-2020. Before the study, the participants signed a written informed consent form. The final analyses were conducted with a sample size of 46 (19 female, 20 male, 7 participants’ gender and age data were not available, for eliminations, see below).

### Smartphone use monitoring

Smartphone use was quantified with an app ( TapCounter, QuantActions AG, Zurich), which collects (i) the timestamp of touchscreen interactions on the smartphone, (ii) the active app, and (iii) the screen lock and unlock times. The app was installed at least a week before the laboratory EEG recording session. The timestamps were collected at millisecond resolution (mean error = 0, standard deviation = 15 milliseconds) [Balerna and Ghosh, 2018]. We leveraged the period before the recording session to determine the commonly used apps based on the number of interactions. In the laboratory, the participants were asked to engage with their top 2 apps in the social and non-social categories (for definitions see: [Westbrook et al., 2021]). Besides this request, they were asked not to view videos or listen to music, and instructed to use their right thumb on the screen.

### Tappigraphy sleep duration

Sleep duration was estimated based on smartphone usage patterns (tappigraphy). This method of estimating sleep duration has been validated previously and contrasted to sleep durations collected through wearable devices [Borger et al., 2019, Massar et al., 2021]. Briefly, the algorithm leverages circadian patterns and the low smartphone activity levels at night to estimate sleep durations. We estimated sleep duration for the night preceding the experiment, as well as the median sleep duration across the seven nights before the experiment.

### EEG data collection sessions, processing, and slow wave identification

EEG data were collected while participants engaged in passive movie watching or smartphone use in a Faraday cage. The Faraday cage still contained Wi-Fi connectivity to enable smartphone use. The EEG collection was performed using a 64-channel EEG cap and the BrainAmp 64-channel DC amplifier (Brain Products GmbH Gilching). The data were sampled at 1000 Hz. The cap contained 62 equidistant Ag-AgCl electrodes distributed on the scalp (EasyCap GmbH, Worthsee) and 2 electrodes on the face to capture eye blinks. To establish a baseline resting state matching the smartphone use in terms of duration, participants first watched a movie (David Attenborough, Africa series, 2013) for ∼60 minutes. To control for somatosensory stimulation, the movie watching condition included unpredictable tactile stimulation to the fingertips, matching the tactile feedback during smartphone use (for details see [Ghosh, 2021]). This was followed by a period of ∼60 minutes during which they interacted with their smartphone (see Supplementary Figure 1 for the duration of the sessions). Participants were offered brief rest periods ( ∼5 minutes) every 10-15 minutes during smartphone use, which were excluded from analysis. During the measurement, the impedances were maintained below 10 kΩ, and any exceptions were eliminated from further analysis (see below). A subset of the participants (14) performed two separate sessions with identical study procedures, conducted approximately one week apart, as part of a larger project. The measurement with the longest smartphone recording was selected for analysis. Due to a technical error, 6 participants had less than 15 minutes of smartphone data and were excluded from further analyses.

We processed the EEG data offline using MATLAB 2023b (MathWorks, Natick) and EEGLAB [Delorme and Makeig, 2004]. The pipeline for slow wave detection included the following procedures commonly used for sleep slow wave detection [Riedner et al., 2007, Bernardi et al., 2015] and is based on an algorithm adapted for wake slow waves [Andrillon et al., 2021]. First, any channels with high impedance (exceeding 10 kΩ) were identified and replaced with interpolated values using neighboring electrodes (spherical interpolation, *pop interp*). Blink-related artifacts were removed using *icablinkmetrics* ([Pontifex et al., 2017]) based on Independent Component Analysis (Infomax). The data were then band-pass filtered between 0.5 and 48 Hz, followed by downsampling to 128 Hz. Data were first average-referenced for artifact detection, then re-referenced to linked mastoids for slow wave analysis. Data were low-pass filtered at 4 Hz to isolate activity within the delta frequency band. Candidate slow waves were identified based on negative half-waves with durations between 0.1 and 1 s. Waves with positive peaks *>* 75 *µ*V or occurring within 1 s of a large-amplitude event (absolute amplitude *>* 150 *µ*V) were discarded. Finally, the 10% of waves with the largest peak-to-peak amplitudes per channel were selected for analysis. After identifying slow waves, we estimated three key features: (i) density, defined as the number of slow waves per minute; (ii) their mean absolute peak-to-peak amplitude; and (iii) their mean downward and upward slopes, calculated as the average change in voltage per second between the start of the wave and the negative peak, and the negative peak and the positive peak, respectively.

### Mass-univariate analysis of slow wave features across rest and smartphone use

To test whether slow wave density fluctuated as a function of time elapsed, we used a hierarchical linear modeling approach spanning all electrodes. At the first level, we fitted a regression model per participant linking the number of slow waves within a 1-minute time window to (i) session type (restful movie watching vs. smartphone use), (ii) time elapsed within session (in min; mean-centered per session), and (iii) their interaction. This allowed us to capture both an overall shift in slow wave density between sessions as well as session-specific time-related changes in slow wave density. At the second level, we tested the *β* coefficients across participants using mass-univariate one-sample t-tests with multiple comparison correction in the form of cluster-based permutation testing (FieldTrip ft timelockstatistics; *α* = 0.05; 1000 randomizations; [Maris and Oostenveld, 2007, Oostenveld et al., 2011]). We repeated this procedure for a reduced model without the interaction term to test for session-invariant time-related changes in slow wave density.

Since slow wave density did not vary significantly as a function of time elapsed within sessions, we proceeded to compare slow wave features between the restful movie watching and smartphone use sessions using a simpler GLM-based approach. To this end, we performed a series of mass-univariate (electrode-wise) paired t-tests for slow wave density, amplitude, and slopes. As before, we performed multiple comparison correction using cluster-based permutation testing (FieldTrip ft timelockstatistics; *α* = 0.05; 1000 randomizations;[Maris and Oostenveld, 2007, Oostenveld et al., 2011]).

### Linking minute-by-minute fluctuations in slow wave features to smartphone behavioral fluctuations

We assessed the relationship between slow wave features and the number of smartphone interactions using two approaches. Note, given the redundancies between amplitudes and upward/downward slopes, we did not include the slopes in the subsequent analysis. First, we used pairwise Spearman correlations: smartphone interaction counts vs. density, and the interaction counts vs. median amplitude. We computed the values within 1-minute time bins, and correlated them with smartphone interaction counts from the corresponding time windows. The correlations were computed for each individual at each electrode, and their significance was assessed via permutation testing in MATLAB. To correct for multiple comparisons, we used the False Discovery Rate (FDR) method with the Benjamini–Yekutieli adjustment (*α* = 0.05) [Benjamini and Yekutieli, 2001]. However, this correlational approach treated each slow wave feature independently and therefore did not assess how the combination of features relates to smartphone use. Therefore, secondly, we performed time-binned multivariate regressions to correlate the number of smartphone interactions with the slow wave features [Pernet et al., 2011]. These regressions were conducted at the individual level for each electrode (LIMO-EEG Level 1 analysis). At the group level, we performed one-sample t-tests on the resulting *β* coefficients (LIMO-EEG Level 2 analysis). To correct for multiple comparisons across electrodes and participants, we applied spatiotemporal clustering (as implemented in LIMO) with 1000 bootstraps and a significance threshold of *α* = 0.05.

### Fine-grained dynamics of wake slow waves

We captured the interval dynamics of wake slow waves using the joint-interval distribution (JID). The JID estimates the probability of observing two consecutive intervals between slow wave off-states: interval K and the following interval K+1. The off-states were timed according to the negative peak of the waveform. Each interval pair (K and K+1) was assigned to a bin in a 2D matrix, and kernel density estimation was used to estimate the joint probability distribution over these bins [Duckrow et al., 2021]. The JIDs were computed separately for each participant, session (restful movie watching and smartphone use), and electrode.

We used non-negative matrix factorization (NNMF) to extract prototypical patterns of slow wave dynamics across the scalp. NNMF is an interpretable dimensionality reduction technique that decomposes a matrix into low-rank parts-based representations [Lee and Seung, 1999]. To determine the optimal number of representations (ranks) for each participant, we performed cross-validation by: (i) randomly masking 20% of the matrix, (ii) running NNMF 50 times for ranks 1–10 with random initializations, (iii) reconstructing the original matrix from the NNMF output, and (iv) selecting the rank with the lowest average reconstruction error.

Since NNMF is non-convex and may produce alternative solutions across runs, we selected a stable solution by using the stable and reproducible NNMF (STAR-NNMF) method, as developed by [Ceolini and Ghosh, 2023]. STAR-NNMF involved running 1000 NNMF repetitions and selecting the decomposition with the highest median pairwise cross-correlation across runs. This process yielded, for each participant, prototypical JIDs (meta-JIDs) and their corresponding spatial distributions across the scalp (meta-locations), indicating where these patterns were most prominent. STAR-NNMF was performed separately for each participant and session.

At the group level, we clustered the meta-JIDs using k-means to identify population-wide patterns. We selected the optimal number of clusters (between 1 to 10) by using the Silhouette method, which measures the similarity within and between clusters. Since K-means is also a non-convex algorithm, we performed 1000 repetitions with randomly initialized values and selected the repetition with the smallest sum of squared distances [Lisboa et al., 2013].

### Linking fine-grained wake slow wave dynamics to smartphone use

To assess the differences in slow wave JIDs between rest and smartphone use, we contrasted the conditions with paired t-tests at each electrode using the LIMO-EEG toolbox [Pernet et al., 2011]. This electrode-wise comparison was chosen because direct comparison of NNMF-derived patterns was not feasible, as the number of low-rank components varied across participants. To control for multiple comparisons, we applied clustering of the two-dimensional JID bins with 1000 bootstraps and an *α* level of 0.05. Second, we measured the relationship between consecutive slow wave intervals and the rate of touchscreen interactions. The rate of interactions was estimated based on the number of touchscreen interactions occurring between the corresponding slow waves, divided by the combined duration spanning the two consecutive intervals (K and K+1). We assigned each rate to a bin in the 2D joint-interval space. For bins containing multiple interval pairs, we summed interaction counts and total durations to calculate a bin-level rate. The rate of smartphone interactions was estimated for each participant and electrode. To analyze the population-level patterns, we estimated the median interaction rates across participants for each electrode. Third, we analyzed the latency to smartphone interactions after slow waves with distinct dynamics. Towards this, we estimated the median time (latency) to the next touchscreen interaction from the end of the second interval (K+1) captured in the JID two-dimensional bin. Finally, we quantified whether the features of consecutive slow waves varied according to their interval dynamics. At each two-dimensional bin of the JID, we estimated the median peak-to-peak amplitude and the median duration of the slow waves.

### Data and code availability

The publication will contain links to the data shared on Dataverse.nl. The data is published 1 month after publication. The code is available on GitHub: https://github.com/CODELABCODELIB/slow_waves_2024.

## Author contributions

A.G. conceived the study. R.K., D.H., and A.G. designed the study. R.K. and D.H. analyzed the data aided by A.G. A.G. drafted the report aided by all authors. R.H. contributed with the methods for slow wave detection. All authors helped edit the manuscript.

## Acknowledgments

The authors would like to thank the student assistants who contributed to the data collection at Leiden University.

## 5 Legends

**Supplementary Figure 1.**
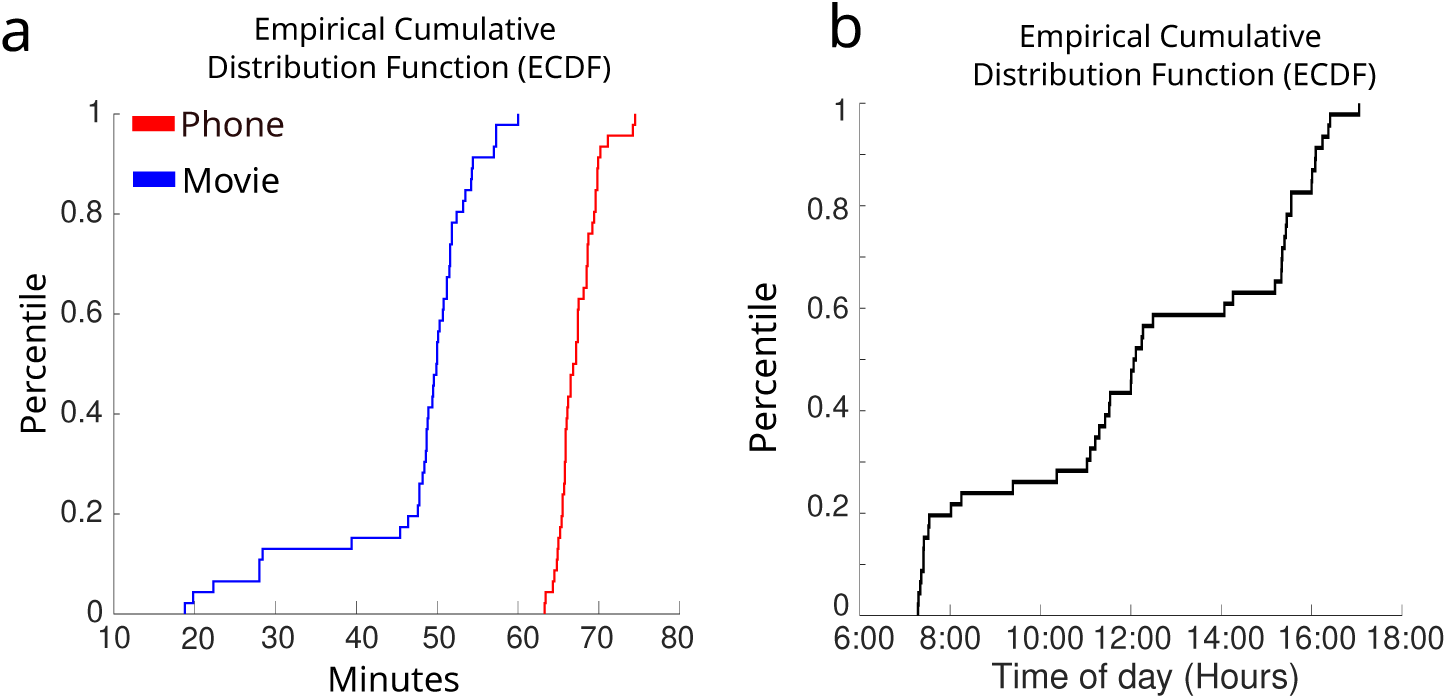
Recording times and session start times. **(a)** Empirical cumulative distribution function (ECDF) of recording times during restful movie watching (blue) and smartphone behavior (red). **(b)** ECDF of the time of day when participants started their measurements.

**Supplementary Figure 2.**
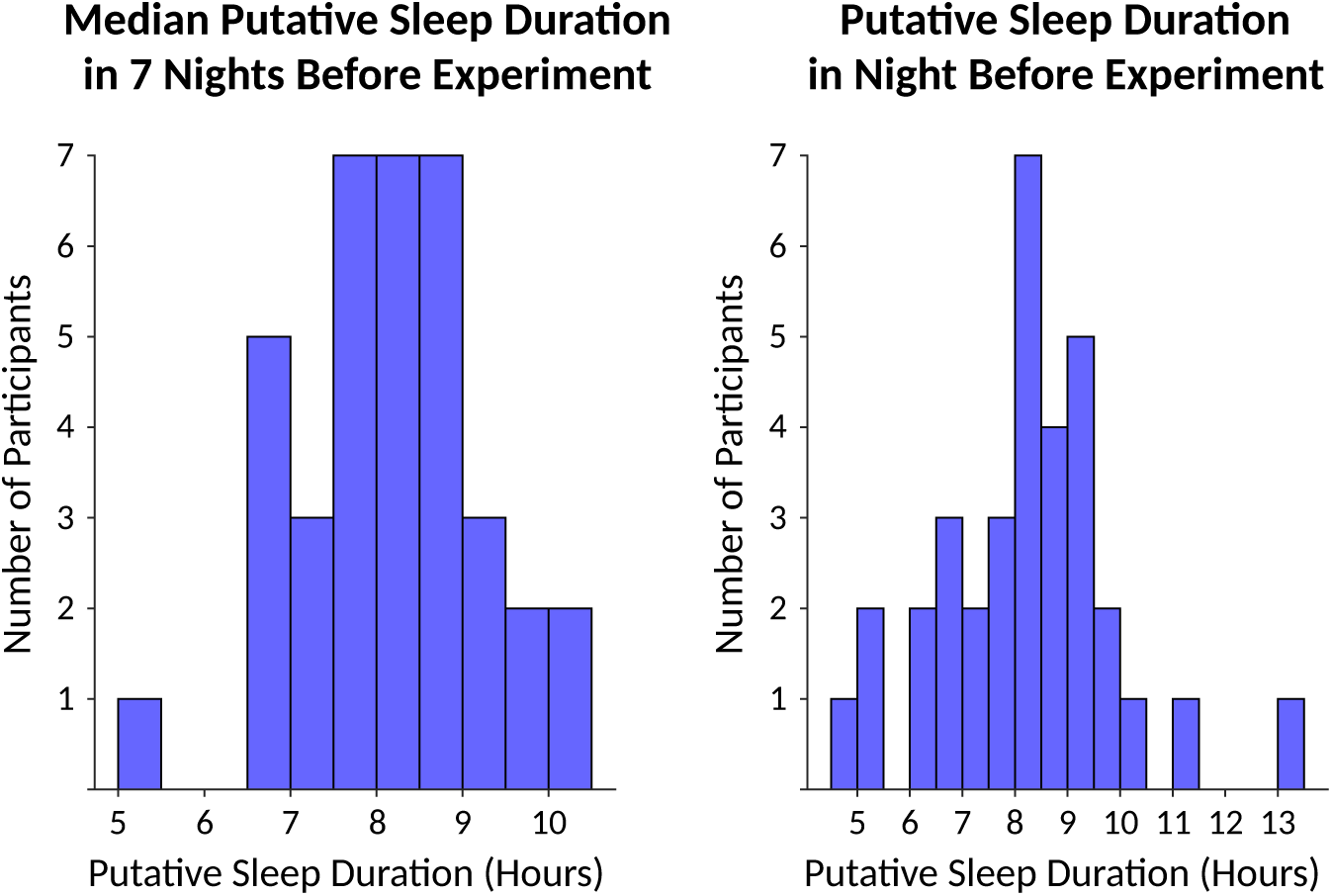
Putative sleep durations prior to the measurement. **(a)** Sleep duration on the night before the measurement, estimated from smartphone usage patterns. The distribution suggests that participants were not experiencing extreme sleep deprivation. **(b)** Median sleep duration across the week before the experiment, also estimated from smartphone data, similarly shows no extreme sleep deprivation. Two participants were excluded from these figures due to missing prior sleep data.

**Supplementary Figure 3.**
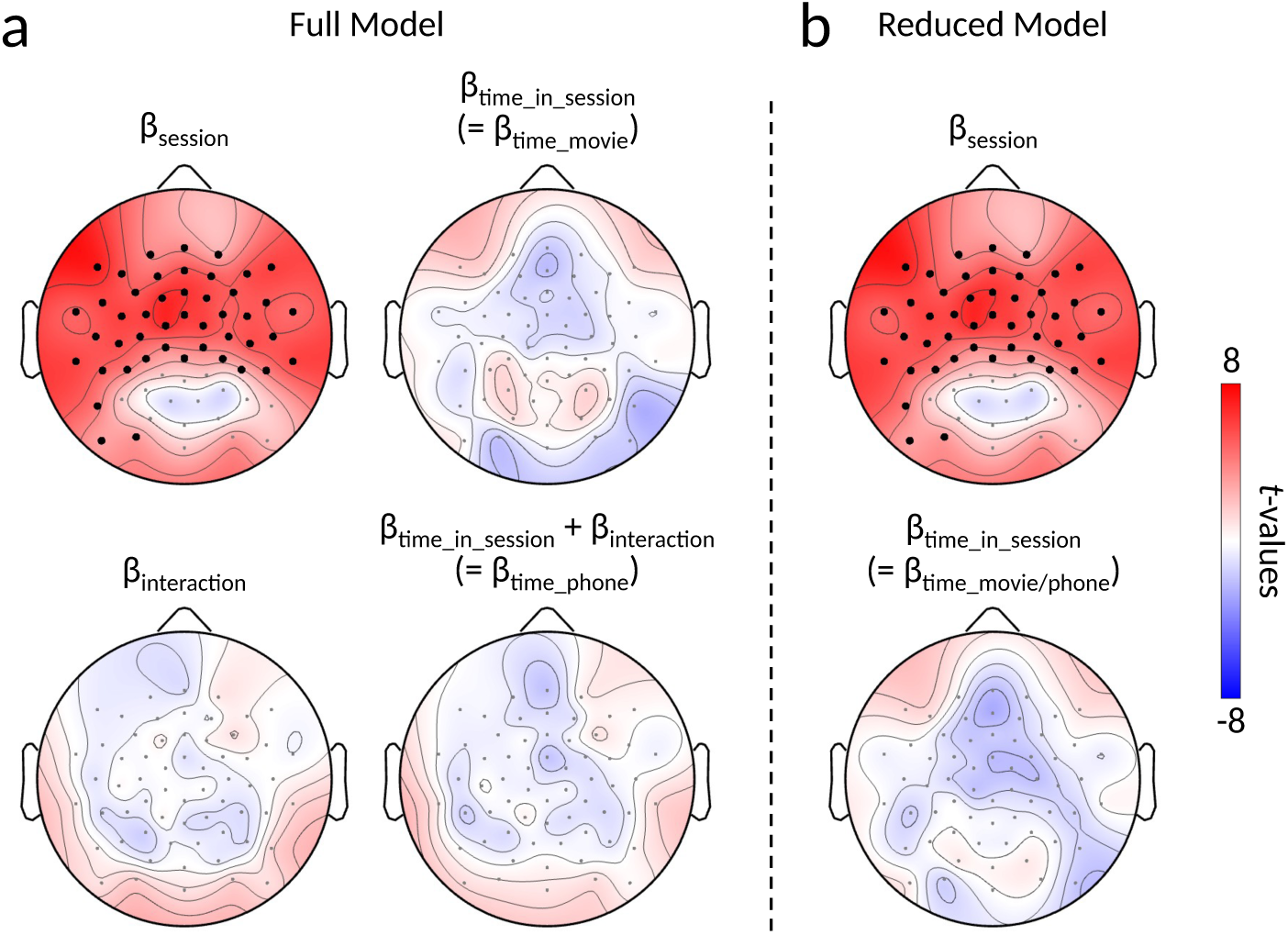
Temporal stability of slow wave occurrence. **(a)** A mass-univariate linear regression was used to test whether slow wave occurrence (number of slow waves per 1-minute bin) changed over time. Topographical plots denote *t* -values for beta coefficients of: (i) session type (restful movie watching or smartphone use), (ii) time elapsed within the sessions (in minutes; mean centered per session), and (iii) the interaction of the two variables. Statistical significance was assessed with multiple comparison correction in the form of cluster-based permutation testing (*α* = 0.05; 1000 randomizations). Only the session type had statistically significant electrodes (black dots), indicating that the number of slow waves changed as a function of behavioral context (smartphone use or rest) but not as a function of time elapsed. **(b)** This analysis was repeated for a reduced model without the interaction term. Again, the effect of time elapsed did not reveal any significant electrode clusters.

**Supplementary Figure 4.**
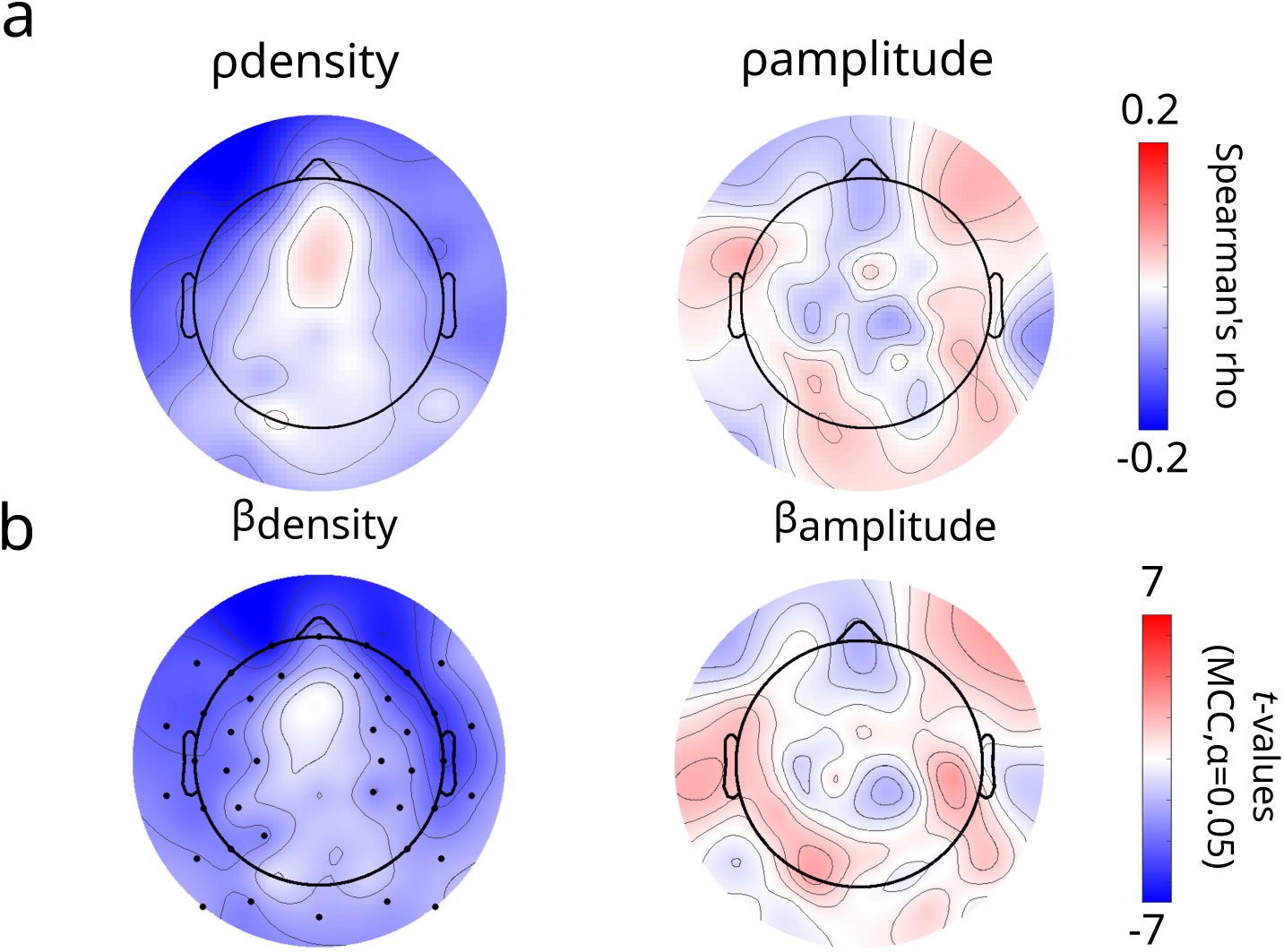
Coarse-grained relationship between slow wave features and smartphone interactions. **(a)** Topographical maps show population-average Spearman’s *ρ* values between the number of smartphone interactions and slow wave density (left) or slow wave amplitude (right), computed within corresponding 1-minute time bins. No reproducible statistically significant correlations were found across the population. **(b)** Hierarchical mass-univariate regressions were performed to test whether slow wave features predict the number of smartphone interactions within corresponding 1-minute time bins. Topographical maps show *t* -values for slow wave density (left) and slow wave amplitude (right). Statistically significant clusters (black dots) emerged primarily over frontotemporal and sensorimotor regions, where higher slow wave density was associated with fewer smartphone interactions. Statistical significance was assessed using spatiotemporal clustering with multiple comparison correction, as implemented in LIMO EEG (*α* = 0.05, 1000 bootstraps)

**Supplementary Figure 5.**
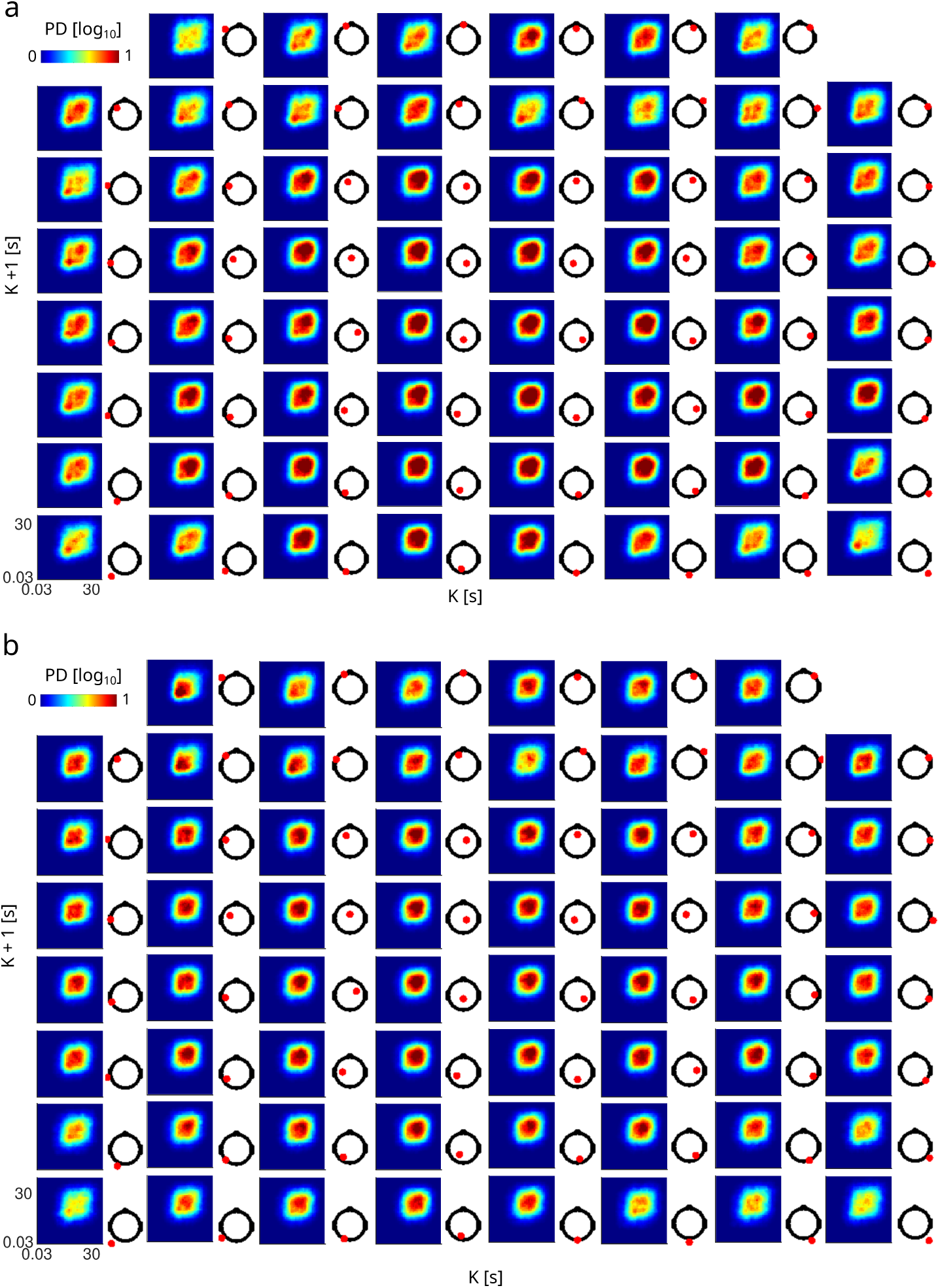
Slow wave joint-interval distributions (JIDs) across the population for all electrodes. **(a)** JIDs during restful movie watching, pooled across participants using the median. Distributions are shown in *log*_10_ scale. Electrode locations are marked with red dots. **(b)** Same as (a), but for JIDs during smartphone behavior. PD = probability density.

**Supplementary Figure 6.**
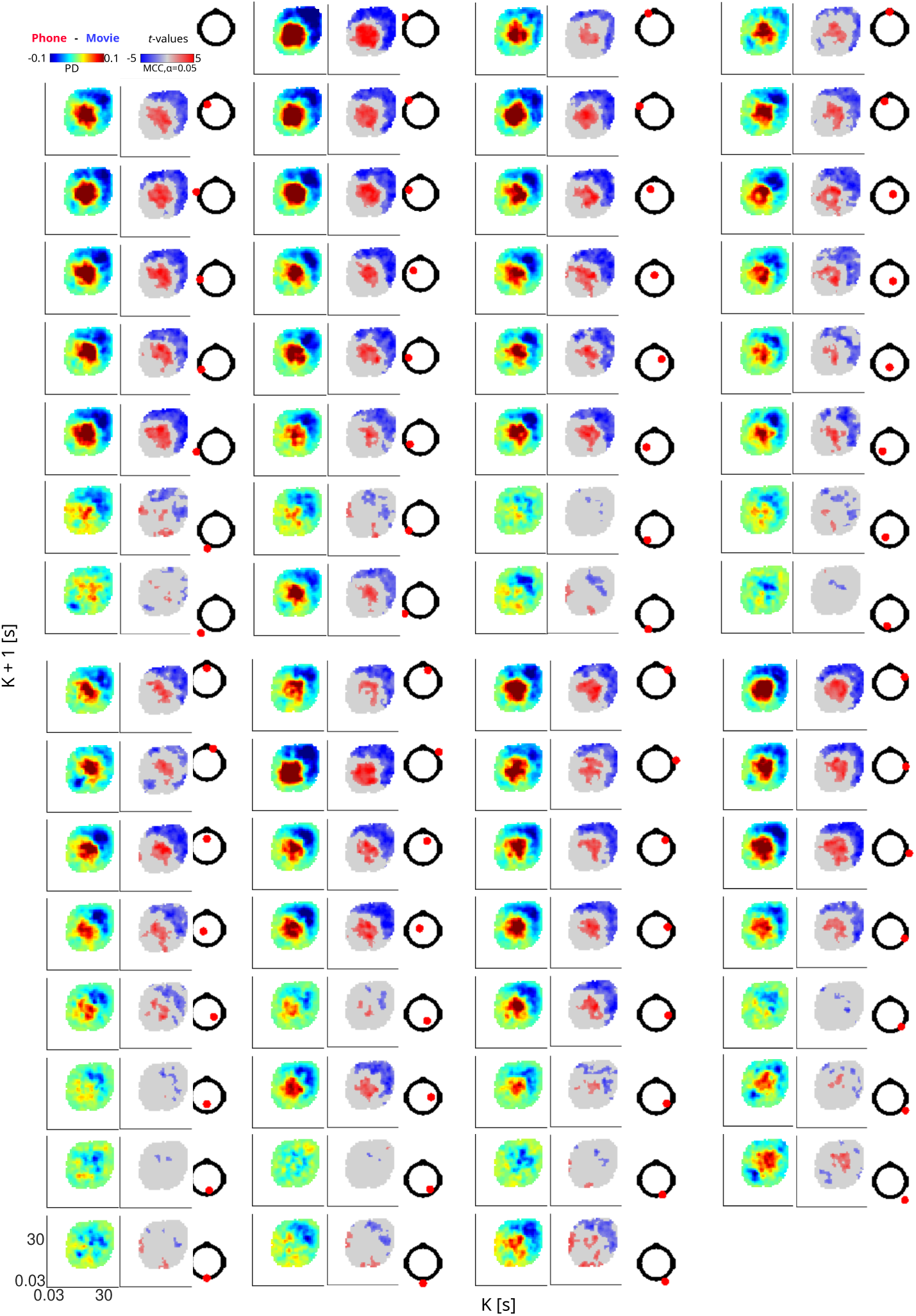
Comparison of slow wave JIDs between movie watching and smartphone behavior across all electrodes. We performed paired t-tests between the two sessions. Mean differences between slow wave JIDs (smartphone behavior – restful movie watching) are shown in the left column. Statistically significant *t* -values are displayed in the middle column. Electrode locations (marked with red dots) are displayed in the right column. Statistical significance was assessed using spatiotemporal clustering with multiple comparison correction, as implemented in LIMO EEG (*α* = 0.05, 1000 bootstraps). Grey indicates no statistically significant clusters, and white indicates that there were no events in the bins. PD = probability density.

**Supplementary Figure 7.**
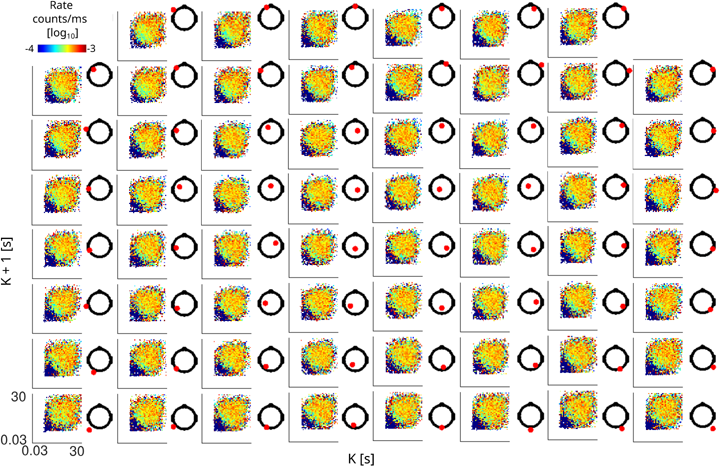
Population-level medians for behavioral rate across slow wave JIDs for all scalp electrodes (red dots denote electrode location). The rate of smartphone interactions was lower during rapid bursts of slow waves and higher during sparser consecutive slow waves. The pattern only marginally varied across the scalp. White areas indicate bins with no detected events.

**Supplementary Figure 8.**
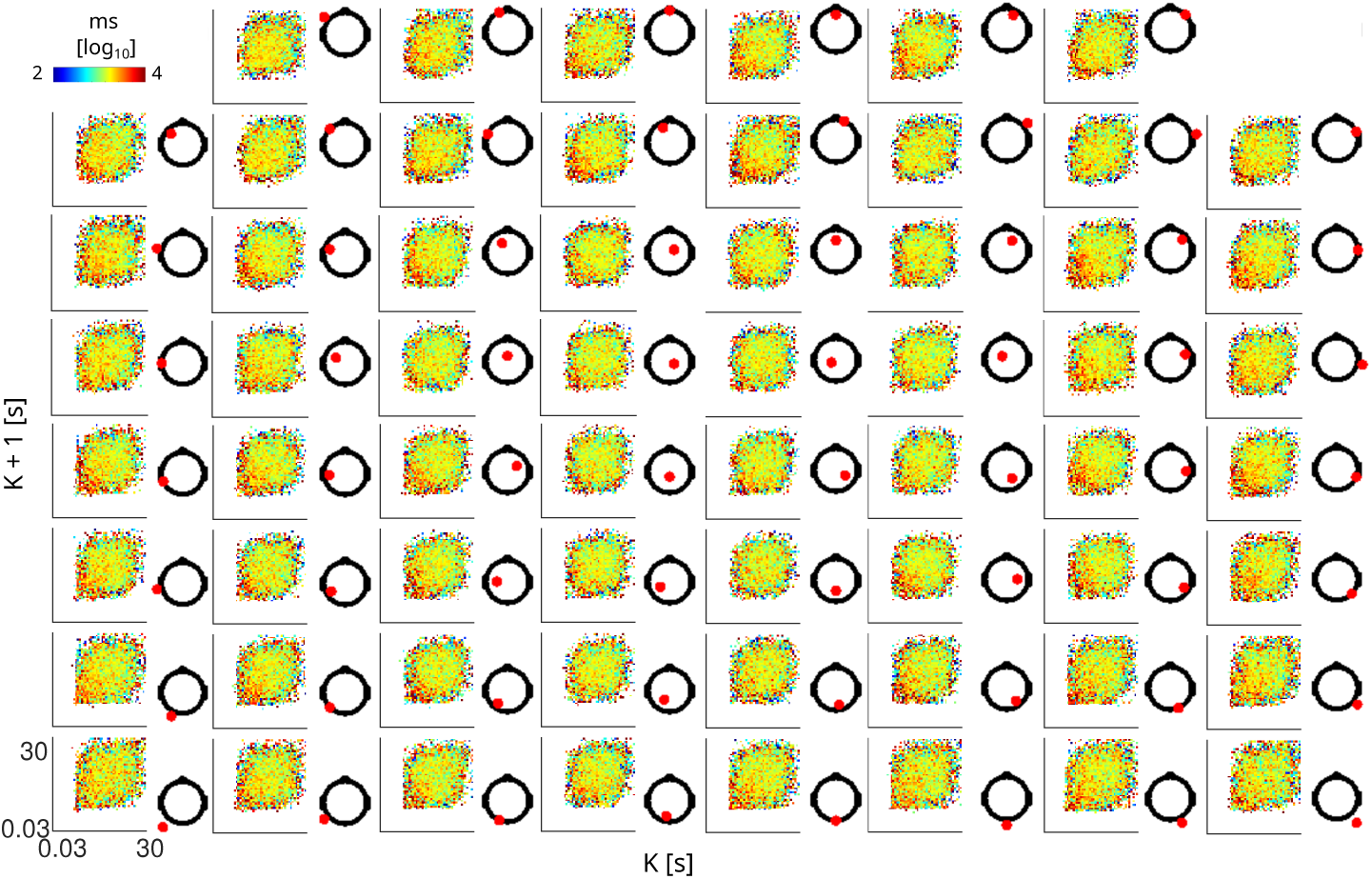
Median latency to the next smartphone interaction following consecutive slow waves. A marginal gradient was observed: the next interaction tends to occur later following rapid consecutive slow waves, and sooner following sparse ones. White areas indicate bins with no detected events.

**Supplementary Figure 9.**
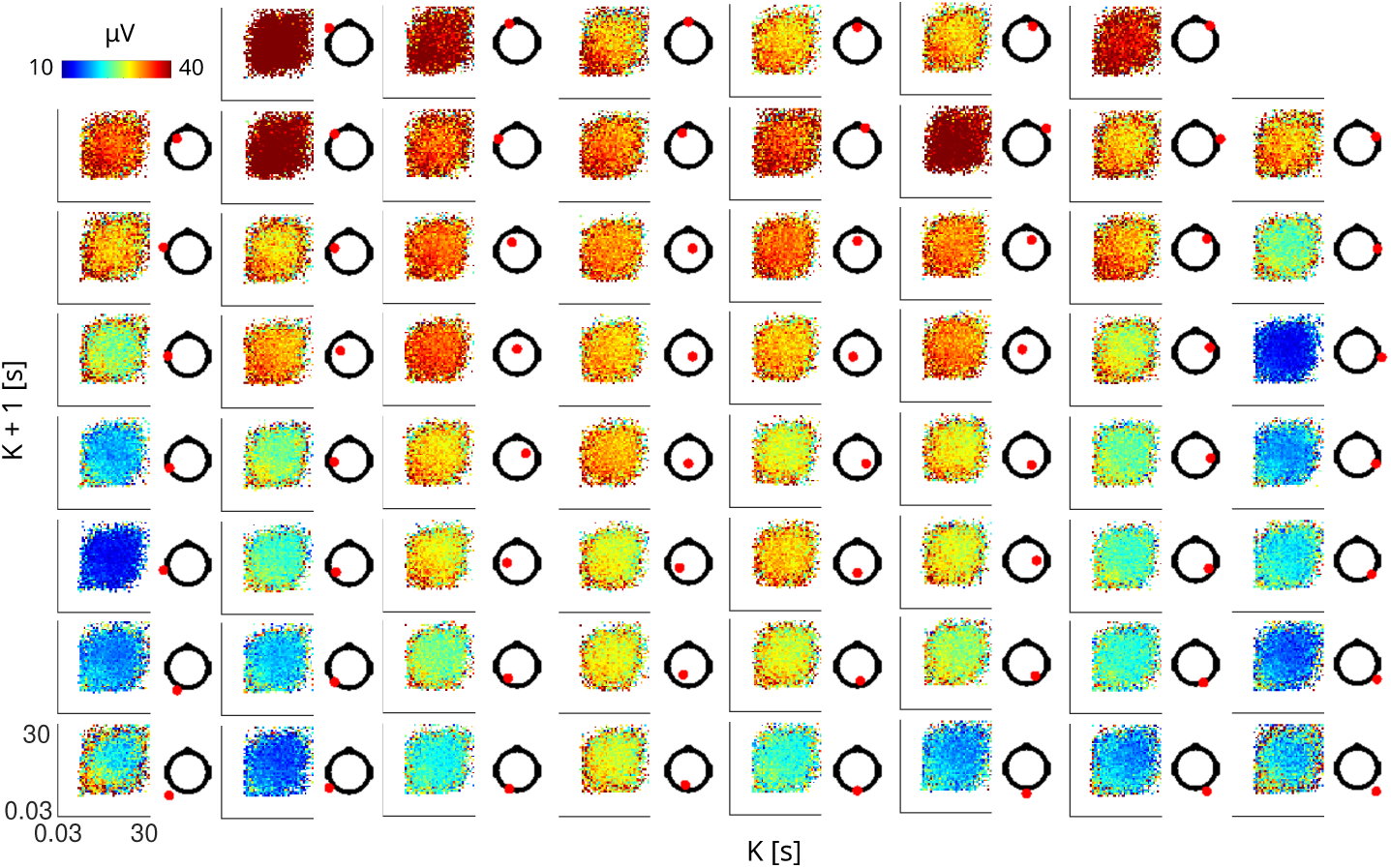
Population-level slow wave amplitudes and slopes projected onto the JID. **(a)** Median peak-to-peak amplitudes were calculated across consecutive slow waves. Amplitudes at each electrode (red dots) were pooled across participants using the median. A gradient along the diagonal showed that the amplitudes decrease as slow waves become sparser. Although this gradient was consistent across electrodes, the average amplitudes varied. White areas indicate bins with no events.

**Supplementary Figure 10.**
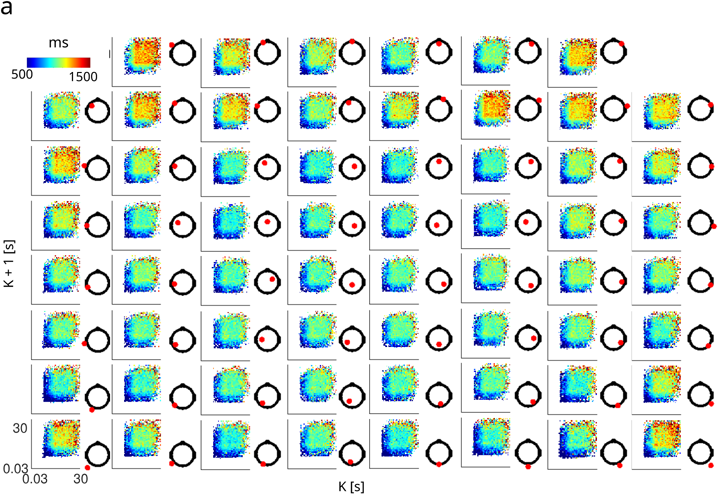
Median duration of slow waves (from onset to offset) occurring across consecutive intervals. For each electrode (red dot), we show the resulting median population-level durations projected onto the JID. A gradient emerged: durations were generally shorter during rapid consecutive slow waves and longer during sparse ones. White areas indicate bins with no events.

